# Integrated Epigenetic and Transcriptomic Profiling Reveals Dynamic Regulatory Networks Driving Retinoblastoma Pathogenesis

**DOI:** 10.1101/2023.05.10.540147

**Authors:** Yihan Zhao, Peng Liu, Lei Hu, Yu Shi, Yan Li, Yanting Wang, Zehua Zeng, Hongwu Du, Binzhi Qian

**Author notes:** These authors contributed equally to this work (Yihan Zhao, Peng Liu). All correspondence should be addressed to, and.

## Abstract

Retinoblastoma (RB), the most aggressive pediatric intraocular malignancy, urgently requires mechanistic insights to overcome limitations of current clinical interventions. Through integrated single-cell multi-omics analysis, we constructed a comprehensive epigenetic-transcriptomic atlas of photoreceptor subpopulations in RB. Pseudotemporal trajectory analysis revealed the differentiation cascade from retinal progenitor cells (RPCs) to malignant cone-like tumor cells (Cone-T), orchestrated by dynamic regulation of tumor suppressors (FEZ1) and oncogenic drivers (CITED2) within the MAPK pathway. Notably, we identified three photoreceptor subtype-specific transcription factors (EBF1, SOX15, NFIL3) exhibiting concordant overexpression and chromatin accessibility. Cell-cell communication analysis uncovered PPIA-BSG ligand-receptor interactions potentially driving tumor progression, while drug screening prioritized nine therapeutic candidates targeting transitional proliferative photoreceptors (P-p). Our findings establish a mechanistic framework for photoreceptor transformation and provide actionable targets for precision therapies.

## 1 Introduction

Retinoblastoma (RB) remains a highly invasive intraocular malignancy primarily affecting infants and young children, with an incidence of approximately 1 in 16,000 and a median age of onset at 18 months[1]. Characterized by biallelic mutations in the RB1 gene within developing retinal cells [2] [3], untreated RB can invade the optic nerve, leading to blindness. The degree of histopathological invasion is a strong predictor of metastasis risk [4] [5]. With advances in medical technology, therapeutic options for retinoblastoma have diversified to include chemotherapy, laser photocoagulation, and thermotherapy [6], enucleation remains unavoidable in the absence of early intervention or in cases with severe complications such as edema, conjunctival cysts, pyogenic granulomas, ptosis, and superior sulcus defects [7].

Current studies on retinal development have proposed three hypotheses regarding the cellular origin of retinoblastoma, which include retinal progenitor cells, cone photoreceptors, and either horizontal cells or Müller glial cells[8]. Recent evidence increasingly supports G2/M cone precursors as the likely cells of origin for RB [9] [10] [11]. However, these hypotheses primarily focus on the origins and developmental progression of retinoblastoma, leaving the underlying mechanisms of invasion, metastasis, and specific regulatory pathways largely undefined. Further investigation is thus crucial to refine and validate these models.

Single-cell RNA sequencing (scRNA-seq) has become an invaluable tool for dissecting gene expression at single-cell resolution, facilitating the exploration of cellular heterogeneity and early embryonic development [12]. Complementarily, single-cell ATAC sequencing (scATAC-seq) analyzes accessible chromatin regions to identify active DNA regulatory elements, proving particularly useful in studies of developmental regulation [13] [14]. To enhance regulatory analysis, the GLUE algorithm [15] explicitly models regulatory interactions, bridging distinct feature spaces across omics layers. Furthermore, algorithms like SIMBA [16] [17] enable the integration of omics-defining features into a shared latent space, providing novel insights into tumor regulatory mechanisms and the identification of potential therapeutic targets.

In this study, we present a single-cell resolution transcriptomic and chromatin accessibility landscape of retinal development in the context of retinoblastoma. Our findings reveal that proliferative photoreceptor cells are developmental precursors to Cone-T cells and represent potential cells of tumor origin. We demonstrate that genes including HSPA1A, RP1, DMD, and transcription factors such as EBF1, SOX15, and NFIL3, play significant regulatory roles during tumorigenesis in these cells. This discovery provides a critical theoretical foundation for developing effective treatment and prevention strategies, marking a significant stride towards individualized therapy.

## 2 Results

### 2.1 The Single-Cell Multi-Omics Atlas of the Retina and Retinoblastoma

To comprehensively characterize cellular heterogeneity in both the normal retina and retinoblastoma, we analyzed scRNA-seq data from 71,746 cells and scATAC-seq data from 85,418 cells. These datasets comprised normal retinal samples collected across nine developmental stages, ranging from 9 to 26 post-conceptional weeks (PCW) (9, 10, 11, 14, 15, 18, 20, 23, 26 weeks)[18], and retinoblastoma patient samples from two time points (4 months and 2 years)[19] (Fig. 1A) Following rigorous quality control and batch effect correction, we identified 17 distinct cell types in the scRNA-seq dataset using canonical retinal cell type markers[20], Amacrine cells (AC, high PAX6 and MALAT1 expression); Bipolar cells (BC, high VSX1 and CA10 expression); Cone photoreceptor cells (Cone, high PDE6H and GNB3 expression); Cone-like tumor cells (Cone-T, high MCM3 and PCNA expression); Fibroblasts (FB, high COL1A1 and COL3A1 expression); Horizontal cells (HC, high STMN2 expression); Lens cells (LC, high LIM2 and CRYAB expression); Melanocytes (MC, high CD44 expression); Microglia (MG, high C1QA and HLA-DRA expression); Müller glia cells (Müller, high VIM expression); Red blood cells (RBC, high HBA2 and HBA1 expression); Retinal ganglion cells (RGC, high ISL1 and NEFL expression); Retinal progenitor cells (RPC, high SOX2 and SFRP2 expression); Retinal pigment epithelial cells (RPE, high PMEL and SERPINF1 expression); Rod photoreceptor cells (Rod, high NRL and NR2E3 expression); Rod precursor cells (Rod-P, high NRL and RCVRN expression); and Vascular endothelial cells (VC, high TM4SF1 and CLDN5 expression) (Fig. 1C-D, sfig. 1A,E)

**Fig. 1.**
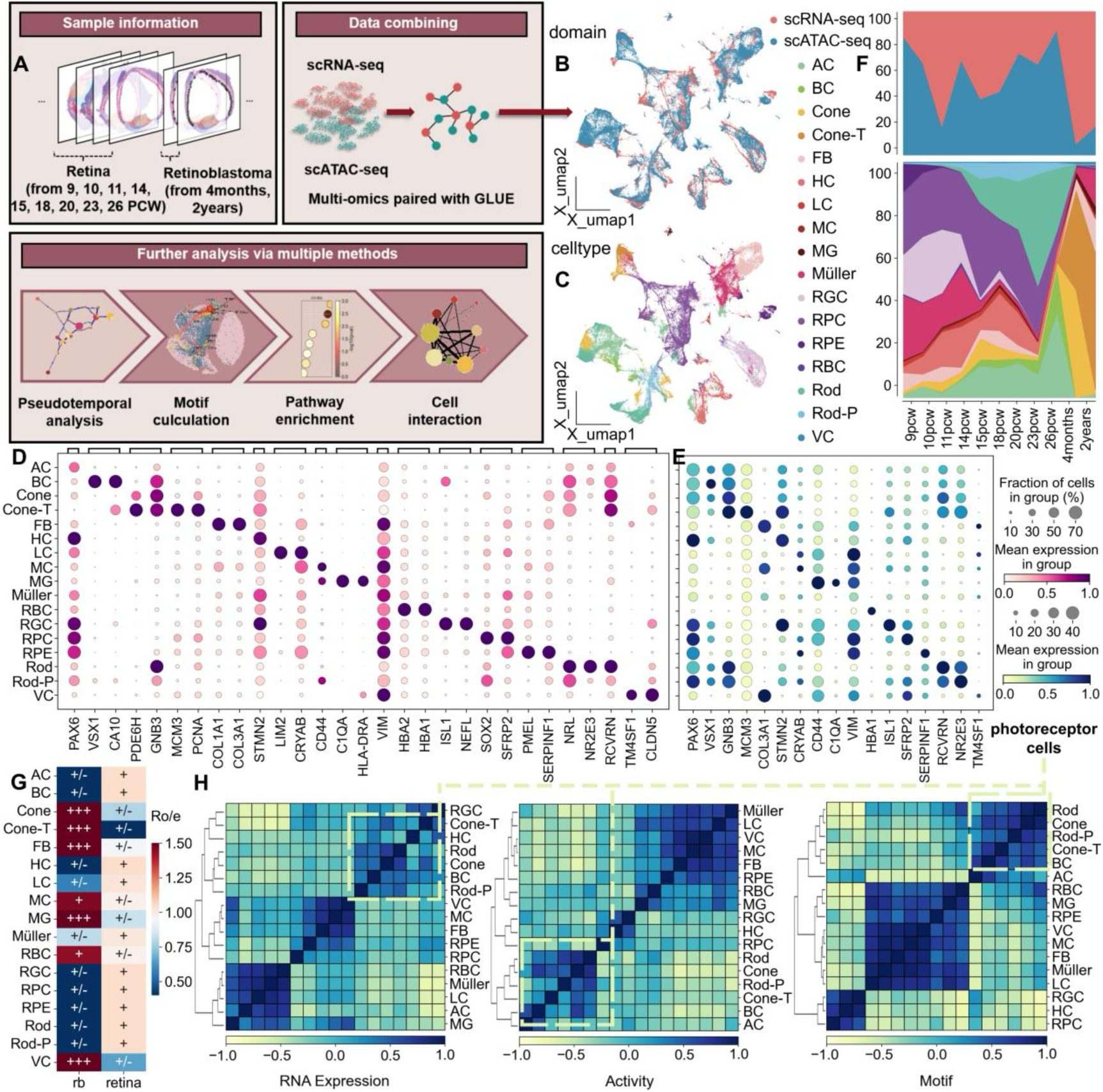
Multi-Omics Landscape of the Retina and Retinoblastoma from scRNA-seq and scATAC-seq Data. **A**, Schematic diagram of sample collection, data combination, and experimental design. **B**, Uniform manifold approximation and projection (UMAP) visualization of integrated multi-omics data with omics layer. **C**, UMAP representation of multi-omics data, with cells categorized by cell type. **D**, Dot plot showing selected marker genes for each cell type. (Fig. 1C). Dot size represents the proportion of cells expressing each gene, and color intensity indicates standardized gene expression levels. **E**, Dot plot showing the activity levels of marker genes in major cell types. Dot size represents the proportion of cells with detectable gene activity, and color intensity represents standardized gene activity values. **F**, Stacked area graph displaying the distribution of domains(top) and celltypes(bottom) across developmental time points (weeks). **G**, The heatmap illustrates the Ro/e values for major cell types across various tissue types. Ro/e > 1.5 indicates that the major cell type is preferred to distribute in the corresponding sample type. More details could be found in the ‘Methods’ section. **H**, Correlation matrix showing RNA expression, ATAC-seq activity, and motif enrichment across different cell types, with color indicating the magnitude of subtype correlation.

To annotate the scATAC-seq profiles, we performed integration of scRNA-seq and scATAC-seq datasets using Graph-linked Unified Embedding (GLUE)[15], and we trained k-nearest neighbors (KNN) model to tranfer the cell type annotation from scRNA-seq to scATAC-seq using embedding of cells(Fig. 1B, sif. 1C). We also calculated gene activity matrices from scATAC-seq profiles and evaluated the activity of scRNA-seq’s marker genes in scATAC-seq to validate the accuracy of our cell type labeling transfers (Fig. 1E, sfig. 1B).

Analysis of cell subtype distribution across tissue types (tumor vs. non-tumor) and developmental stages revealed that BC and HC were exclusively present in normal samples, while Cone-T cells were specific to retinoblastoma samples. Furthermore, Cone cell proportions were markedly increased, whereas RPC proportions were notably reduced in retinoblastoma samples (Fig. 1F-G, sfig. 1D).

Finally, we analyzed correlations between retinal and retinoblastoma cell types based on RNA expression profiles, gene activity levels, and transcription factor binding motifs. Notably, five photoreceptor-associated cell subtypes (RPC, Rod-P, Rod, Cone, and Cone-T) exhibited strong correlations (scores > 0.8) across all three matrices, suggesting that photoreceptor cells possess a regulatory modularity distinct from other retinal cell types (Fig. 1H).

### 2.2 Transcriptional Dynamics in the Development of Tumor and Non-Tumor Photoreceptor Cells

During postnatal development, Rb is expressed in RPC and differentiating Rod [21]. To analyze transcriptional changes in photoreceptor cells from normal retinas versus tumor tissues following retinal degeneration, we investigated the transcriptional dynamics of photoreceptor development in both contexts. We incorporated scRNA-seq data from four additional invasive and non-invasive retinoblastoma samples to broaden the representation of photoreceptor development [22] (sFig. 2A). We extracted photoreceptor cells from the retinal and retinoblastoma scRNA-seq datasets, yielding a total of 30,072 and 20,843 cells, respectively. Using scVI [23] to correct for batch effects, we re-annotated the subtypes of retinal photoreceptor cells (Fig. 2A). Based on subtype-specific marker genes, we identified seven major subtypes: cone precursor cells (CPC: RXRG, PDE6H); cone photoreceptor cells (Cone: PDE6H, GNAT2); proliferative photoreceptor cells (P-p: APOLD1, CENPF); retinal progenitor cells (RPC: CCND1, SFRP2); retinal stem cells (RSC: RBP1); rod photoreceptor cells (Rod: GNAT1, NR2E3); and rod precursor cells (Rod-P: CRX, RCVRN) (sFig. 2B). Within retinoblastoma samples, we similarly identified nine photoreceptor subtypes, including tumor-like cone cells (Cone-T: MCM3) and intermediate cells (Middle: EGFLAM). T-cell receptor-like photoreceptor cells (TCR: ZFAS1, USPL1) were also identified, with other subtypes sharing similarities to those in the retina, with the exception of CPC and RPC.

**Fig. 2.**
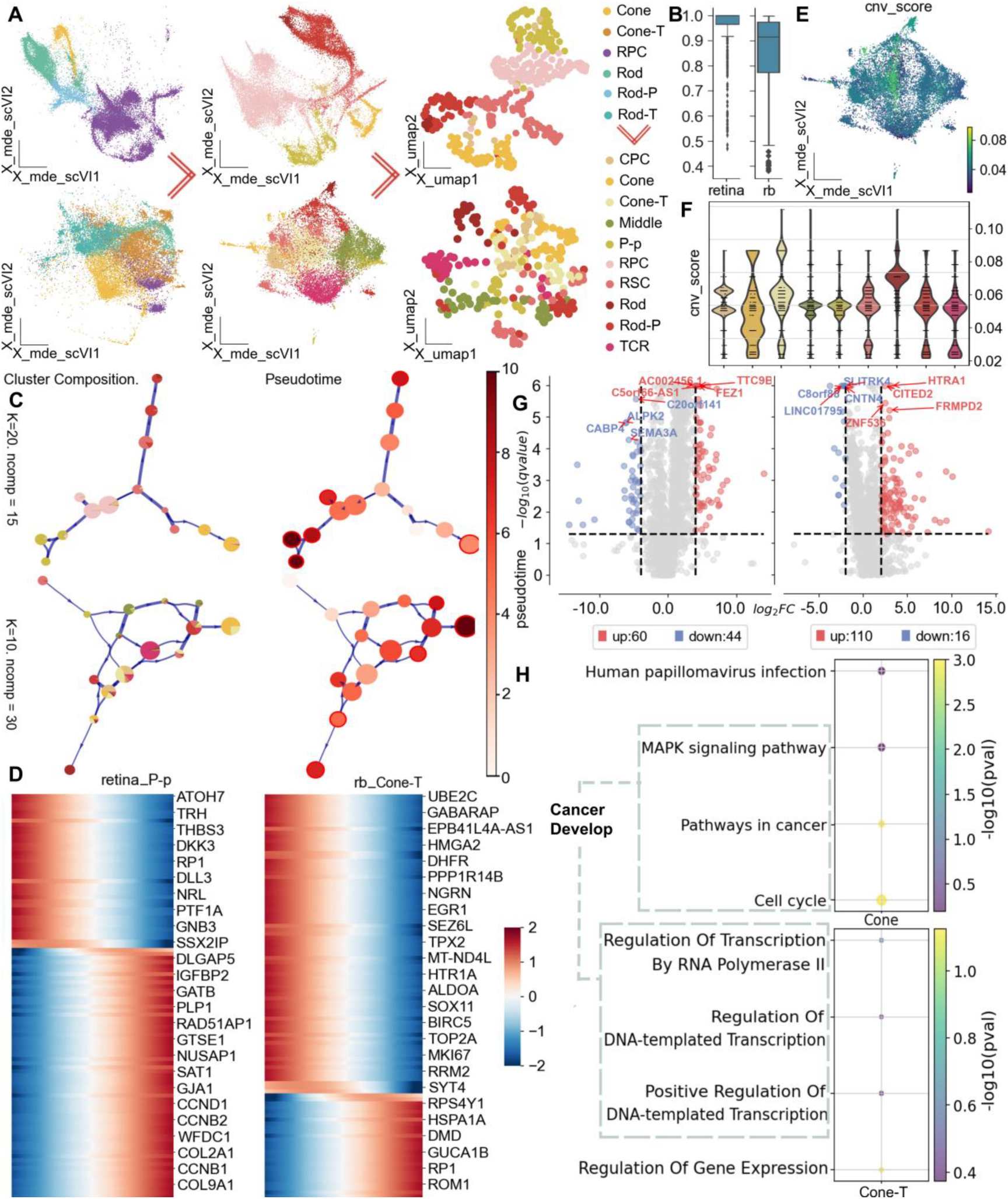
Transcriptional Dynamics Atlas of Retina and Retinoblastoma. **A**, UMAP visualizations showing the extraction of photoreceptor cells (left), projection clustering, re-annotated mapping (middle), and metacell computation (right) for retina (top) and retinoblastoma (rb) (bottom) samples.. **B**, Purity of cell types within metacells. **C**, Pseudotemporal trajectories of photoreceptor cells in the retina (top) and rb (bottom): the left panel displays differentiation trajectories based on subtypes nodes, while the right panel illustrates the trajectories colored by pseudotime. Red circles mark potential differentiation endpoints. **D**, Heatmap of key density driver genes along the developmental trajectories of photoreceptor cells in the retina(P-p) (left) and rb(Cone-T) (right). More details could be found in the ‘Methods’ section. **E-F**, UMAP and violin plots showing copy number variations (CNVs) in different photoreceptor subtypes within rb. **G**, Volcano plots illustrating differential gene expression between P-p and Cone in rb (top), and between Cone and Cone-T cells (bottom). **H**, Enrichment analysis of biological pathways for upregulated genes: visualization of cancer-related pathways in Cone (top) and biological process enrichment for upregulated genes in Cone-T(bottom).

To enhance trajectory inference for photoreceptor cells, we employed the SEACells [24] algorithm to compute metacells and selected 300 and 200 metacells from the retina and retinoblastoma datasets, respectively, at a 100:1 cell ratio (Fig. 2A, sFig. 2C). All metacells exhibited high purity for their respective cell subtypes (Fig. 2B). We then utilized the pyVIA algorithm [25] to infer scalable cell differentiation trajectories and identify lineage fates in photoreceptor cells from both the retina and retinoblastoma samples. In the retina, the developmental trajectory initiates with RSCs and proceeds along three differentiation pathways: 1. RSC → RPC → P-p; 2. RSC → Rod-P → Rod; and 3. RSC → CPC → Cone. These pathways converge at three endpoints: P-p, Rod, and Cone, consistent with known developmental trajectories (Fig. 2C, sFig. 4A). In retinoblastoma, the trajectory similarly begins with RSCs, differentiating into P-p, which further develops into Cone/Cone-T and Rod subtypes. Thus, the trajectory diverges into two primary paths: 1. RSC → P-p → Cone and 2. RSC → P-p → Rod (Fig. 2C, sFig. 4A). Analysis of developmental trajectories and a fitted ridge regression model (sFig. 2D) revealed that P-p serves as both the terminal stage in normal photoreceptor development and the starting point for malignant differentiation, highlighting their pivotal role in retinoblastoma onset and the emergence of malignant Cone-T cells.

To identify key genes associated with the developmental trajectory of photoreceptor cells, we used the Mellon algorithm [26] to compute gene density scores and their expression trends along pseudotime, thereby revealing driving genes in development (Fig. 2D, sfig. 3B-E, Methods). Notably, CCND1 (Cyclin D1), a driver gene, is crucial for regulating the RPC cell cycle and retinal tissue formation, and its regulatory functions within the retinoblastoma pathway are well-established [27]. In retinoblastoma, we identified driving genes such as HSPA1A [28], RP1 [29], and DMD [30], all recognized as critical factors in cancer progression. Furthermore, the gene ROM1 [31], a predictor of retinoblastoma progression, is significantly upregulated in Cone-T cells, suggesting its role in tumor initiation and exacerbation (Fig. 2D, sfig. 3E).

**Fig. 3.**
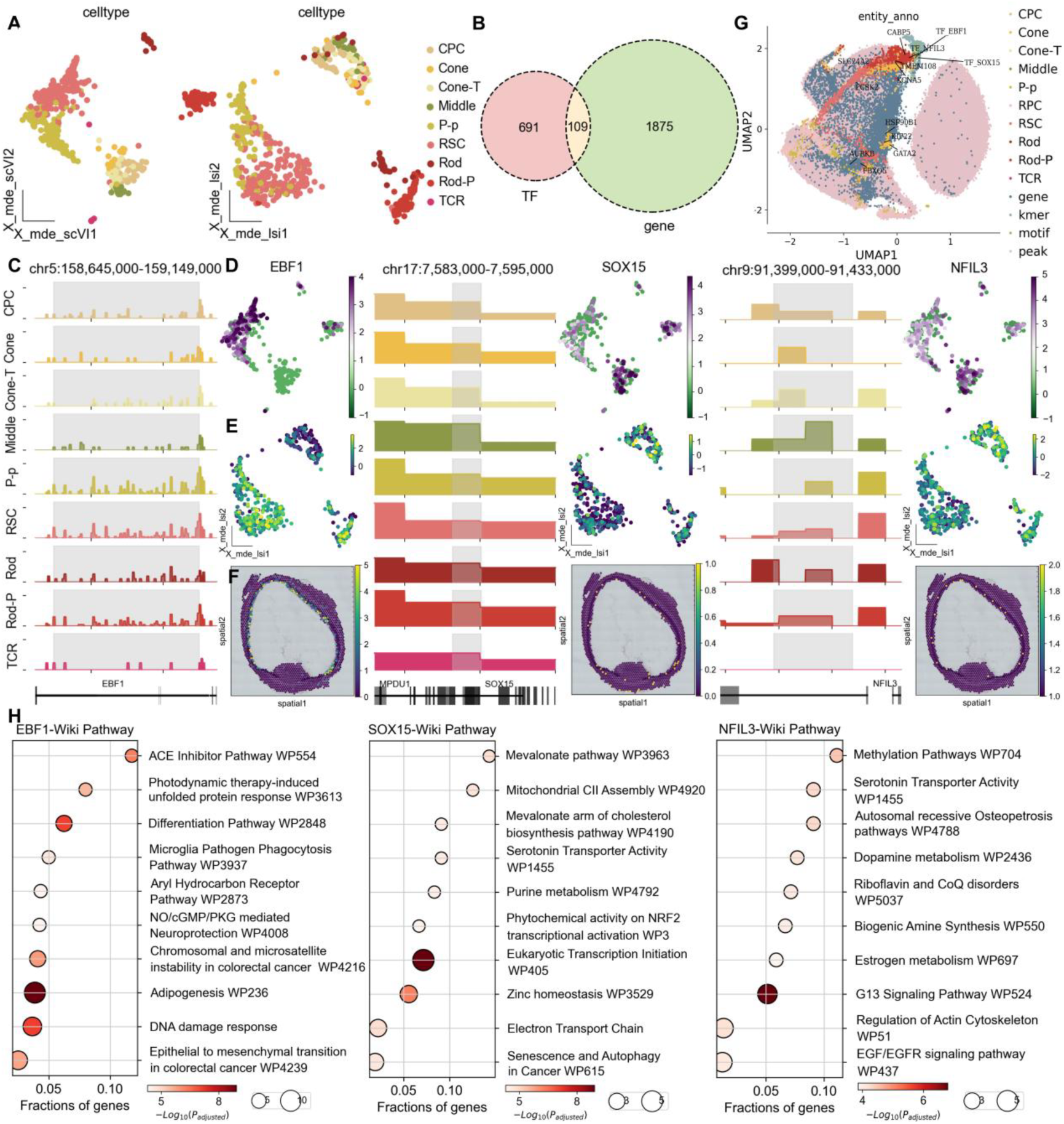
Key Transcription Factor Regulatory Landscape of Retinal and Retinoblastoma Development and Differentiation. **A**, UMAP showing scRNA-seq data (left) and scATAC-seq data (right) of rb, colored by celltypes. **B**, Venn diagrams showing the number of transcription factors (TF) and highly variable genes, as well as their overlaps. **C**, Genomic accessibility tracks of EBF1(left), SOX15(middle), and NFIL3(right) across different rb photoreceptor celltypes. **D**, UMAP showing expression levels of the corresponding genes for the three transcription factors. **E**, UMAP showing the activity scores of three transcription factors. **F**, Spatially resolved expression of three transcription factors in the 13 post-conceptional week (PCW) human retina section. **G**, UMAP plots showing multi-omics data of retinal and retinoblastoma photoreceptor cells, colored by cell subtypes, gene, K-mers, motif, and peak information. Locations of ten temporally regulated genes and three key transcription factors are marked. **H**,Pathway enrichment analysis of the target genes for transcription factors EBF1 (left), SOX15 (middle), and NFIL3 (right) using WikiPathways.

**Fig. 4.**
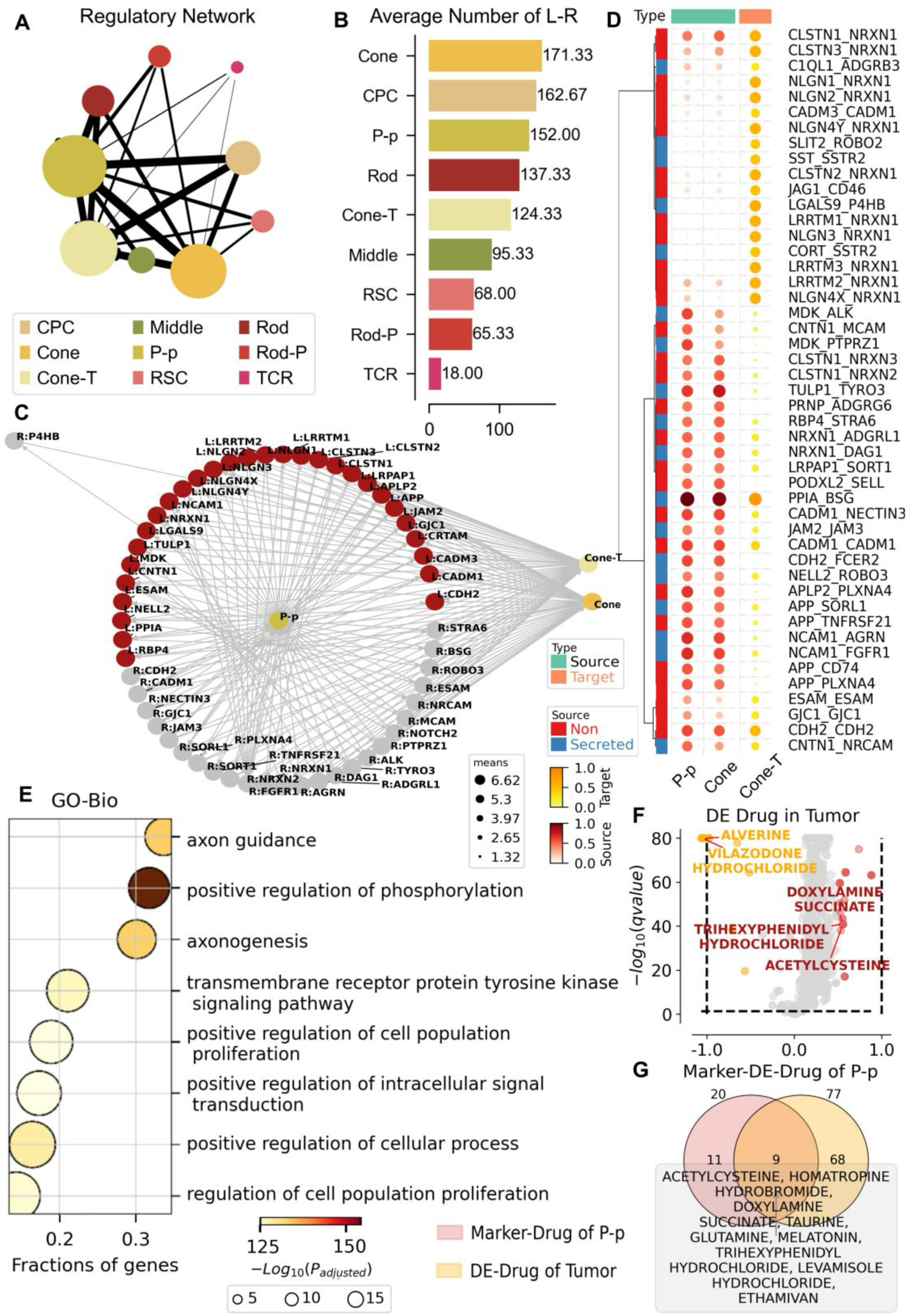
Photoreceptor cell-cell interactions and targeted drug analysis. **A**, Diagram of the intercellular regulatory network. Different colors indicate different cell types, and the thickness of the thread indicates the strength of the connection. **B**, Plot of the average number of ligands and receptors for different cell types. Different colors indicate different cell types, and numbers indicate the average number of ligands for different cell types. **C**, P-p, Cone, Cone-T ligand receptor interaction diagram. The red ones are ligands, the gray ones are the receptors, and the wires indicate the existence of interactions. **D**, Heat map of P-p, Cone, Cone-T ligand receptor interactions. P-p and Cone are the Source, Cone-T is the Target, and the shade of the dot color indicates the intensity of the interaction. **E**, Go enrichment results of highly significant ligand receptors for P-p, Cone, and Cone-T ligands. **F**, Differential drug analysis of cancer cells relative to normal cells. **G**, Intersection of drugs targeting P-p cell types and differential drugs for cancer cells relative to normal cells obtained in Fig 4.F.

To further validate the deterioration trajectory from P-p to Cone-T, we utilized microglia as a reference to assess copy number variation (CNV) [32] scores for all cells in the samples. Our analysis indicated that Cone-T cells exhibited the highest CNV scores, reflecting the greatest malignancy (Fig. 2E-F, sFig. 4A-B). Consequently, we concluded that the trajectory of tumor progression and deterioration proceeds from P-p to Cone and subsequently to Cone-T. By comparing and selecting differentially expressed genes across the differentiation processes of the three subtypes, we observed that during this transformation, the expression of tumor suppressor genes such as FEZ1 [33] and proliferation-associated genes such as HTRA1[34] and CITED2[35] [36] was upregulated (Fig. 2G). In summary, during the differentiation from P-p to Cone, the upregulation of tumor suppressor genes may modulate cell proliferation and the rate of malignant transformation. Conversely, the upregulation of oncogenes during the conversion from Cone to Cone-T accelerates cell proliferation and differentiation, ultimately promoting tumor development.

Further investigation into the biological functions of these two groups of differentially expressed genes was conducted through pathway enrichment analysis. (Fig. 2H). The analysis revealed that highly expressed genes in Cone-T cells were enriched in pathways directly related to cancer cell proliferation and differentiation, including pathways in cancer, cell cycle, and the MAPK signaling pathway [37] [38][39], further substantiatingthe link between differentiation from P-p to cone and the occurrence and progression of retinoblastoma. Additionally, Gene Ontology (GO) enrichment analysis revealed significant enrichment in the biological process of gene expression regulation and the molecular function of cis-regulatory regions/RNA polymerase II transcription regulatory region sequence-specific DNA binding, suggesting that the expression and regulatory functions of relevant genes are disrupted during the development from Cone to Cone-T.

### 2.3 Analysis the regulation of transcription factors in key nodes of tumor development

To identify the transcription regulatory module involved in the development of photoreceptor cells, we paired the scRNA-seq and scATAC-seq profiles using GLUE embedding of cells. Among these, the 729 photoreceptor cells sequenced by scRNA-seq and the 729 sequenced by scATAC-seq are regarded as the same cells, allowing us to indirectly obtain information on two histological layers of a cell (Fig. 3A and sFig. 5A-B). We then inferred transcription factor-associated accessibility using the chromVAR algorithm [40] and identified the top ten transcription factors with the highest activity for the P-p, Cone, and Cone-T subtypes, suggesting that cell differentiation is linked to the regulation of these highly active transcription factors (Supplementary Table1).

To pinpoint subtype-specific regulatory factors, we first selected transcription factors that overlap with highly variable genes (Fig. 3B) and then narrowed it down to the top 50 based on their activity levels. By integrating data from four perspectives—transcription factor activity, gene expression, chromatin accessibility, and gene expression from 13 post-conception week (pcw) retinal tissue sections—we identified three highly specific transcription factors: EBF1, SOX15, and NFIL3 (Fig. 3C-F, sFig. 6A-B), corresponding to the P-p, Cone, and Cone-T subtypes, respectively. We propose that the subtype specificity and regulatory functions of these transcription factors are significant and closely linked to the developmental processes of photoreceptors.

To further refine the regulatory network and determine the target genes of these transcription factors, we employed the SIMBA algorithm [41] to integrate gene data, open chromatin peak information, transcription factor motifs, and k-mer (short DNA fragments derived from sequencing reads) information into a unified graph embedding. After obtaining the SIMBA embedding features, we calculated the unified neighborhood graph for the cells and performed dimensionality reduction visualization. We visualized the locations of the ten inferred fate genes alongside the three specific regulatory transcription factors (Fig. 3G).

Based on spatial positional relationships, we calculated the potential target genes regulated by the three transcription factors (TFs). In the SIMBA spatial map, genes located closer to the TFs are regarded as potential target genes. By calculating a composite score, we identified a total of 6, 50, and 33 potential target genes, respectively (Supplementary Tables2), which were utilized for subsequent enrichment analysis (Fig. 3G). Enrichment analysis of these target genes using WikiPathways (metabolic pathways) revealed that the target genes corresponding to TF_EBF1 are enriched in pathways related to the Cell Differentiation-Index/Differentiation Pathway and NO/cGMP/PKG-mediated neuroprotection, while those corresponding to TF_SOX15 are enriched in the Mevalonate pathway (WP3963) [42] and Eukaryotic Transcription Initiation (WP405) pathways (Fig. 3H). The target genes corresponding to TF_NFIL3 are enriched in the EGF/EGFR signaling pathway [43]. According to the enrichment results, TF_EBF1 primarily mediates the differentiation of photoreceptor cells and the protection of the optic nerve during the precancerous phase, whereas the enrichment pathways of the target genes corresponding to TF_SOX15 and NFIL3 are associated with cancer cell proliferation and metastasis, indicating their regulatory roles in tumorigenesis.

### 2.4 The analysis of intercellular interactions among photoreceptor cells in retinoblastoma

To further explore the cell-to-cell communication during the P-p to Cone tumorigenesis trajectory, we used CellPhoneDB to reveal the rich intercellular communication between photoreceptor cells(Fig 4.A), which also provided a premise for us to further study the deterioration trajectory. As we expected, P-p, Cone, and Cone-T cells all have many receptors and ligands(Fig 4.B), which also indicates that there is a lot of intercellular communication in this deterioration process.

Furthermore, in order to explore the effects of P-p and Cone on tumor formation, we used P-p and Cone as sources and Cone-T as targets, focusing on ligand expression in the source and receptor expression in the target. Interestingly, we found a pair of ligand receptors that were highly expressed in P-p, Cone, and Cone-T: PPIA (Cyclophilin A) and BSG (CD147). (Fig 4.C-D)The interaction between this pair of ligand receptors has been shown to play a vital role in the progression of various diseases such as inflammatory diseases, coronavirus infection, and cancer by activating CD147-mediated intracellular downstream signaling pathways, and is associated with poor prognosis in cancer patients[44][45], suggesting that the formation of retinoblastoma may also be related to this interaction.

We used omicverse to extract highly significant ligand-receptor pairs from these ligand-receptor pairs and performed functional enrichment (Fig 4.E), revealing that positive regulation of phosphorylation may play an important role in the progression of retinoblastoma. role, as mentioned in other cancers[46].

Next, we further conducted drug screening for P-p. Due to the transitional role of P-p in the carcinogenesis of RB, we calculated the drug score for RB (Fig 4.F) and intersected the drugs with high specificity scores with the cell-specific drugs of P-p. Finally, we obtained nine possible potential drugs targeting P-p (Fig 4.G), hoping to provide some suggestions for inhibiting the RB process from a molecular mechanism perspective.

## Discussion

Retinoblastoma, the most prevalent intraocular tumor in children, accounts for 11% of neonatal cancers [47]. Current retinoblastoma (RB) treatment paradigms are progressively evolving toward advanced personalized therapies [48], aimed at preserving vision and mitigating treatment-related risks[49] [50]. Therefore, a comprehensive understanding of RB developmental trajectories and transcriptional regulatory mechanisms is paramount for identifying novel therapeutic targets and designing effective treatment strategies.

This study elucidates the developmental and deteriorative trajectory of photoreceptor cells as RSC → RPC → P-p → Cone → Cone-T, wherein RSCs represent the progenitor cells of retinal photoreceptors[51]. While the association of Cone and Cone-T cells with RB origin and development is well-documented [9] [10], we independently discovered the transitional role of P-p cells, an intermediate developmental state between normal and malignant retinal photoreceptors. This finding further clarifies the process of malignant cone cell generation and offers a novel perspective for preventative RB treatments.

Furthermore, we identified driver genes HSPA1A, RP1, and DMD along the photoreceptor developmental trajectory, and significantly upregulated regulatory genes FEZ1, HTRA1, and CITED2 in Cone and Cone-T cells. Studies have demonstrated that FEZ1 can inhibit cancer cell growth, with its expression inversely correlated with abnormal cell proliferation [33]. CITED2 plays a critical role in fundamental cellular processes including proliferation, differentiation, and migration [35]. Consistent with these findings, our screening of developmental regulatory transcription factors identified subtype-specific transcription factors EBF1, SOX15, and NFIL3 for P-p, Cone, and Cone-T cells, respectively. The literature indicates that EBF1 can suppress transcriptional reactivation of the cancer-associated telomerase catalytic subunit (TERT) at both genomic and epigenomic levels[52],, while SOX15 promotes AOC1 expression and reduces cancer cell proliferation and migration; both transcription factors function as tumor suppressors, impeding cancer development [53]. Conversely, NFIL3 is reported to be highly expressed in various cancers associated with poor prognosis and can inhibit apoptosis induction in cancer cells [54]. In retinoblastoma, the significant expression of EBF1 and SOX15 at early tumor photoreceptor developmental stages suggests enhanced tumor suppressor gene regulation, thereby inhibiting early-stage cancer cell proliferation. Subsequently, in Cone-T cells, the upregulation of transcription factors such as NFIL3 further modulates cone cell differentiation towards malignancy.

Research confirms that CITED2 can interact with other transcription factors and cofactors to promote cell proliferation [36], and TF_NFIL3 can bind and inhibit pro-apoptotic genes (such as TRAIL)[54]. Both factors are specifically regulated in Cone-T cells and favor tumor proliferation. We hypothesize a potential synergistic interaction between CITED2 and NFIL3, wherein CITED2 might enhance tumor cell survival by upregulating NFIL3 expression or activity, consequently suppressing the expression of pro-apoptotic genes like TRAIL.

A limitation of our study is the relatively small sample size. Therefore, our findings are preliminary and require further validation in larger, independent patient cohorts. Additionally, the absence of spatial transcriptomic data from retinoblastoma patients limited our ability to capture spatial information regarding tumor cell heterogeneity. To address this, future studies could utilize engineered mouse models and spatial transcriptomics techniques to further investigate the role of P-p cells in photoreceptor conversion during retinoblastoma development. Furthermore, functional experiments are warranted to validate the identified therapeutic targets, confirming their authenticity and reliability.

In conclusion, this study, leveraging a dual-omics framework of scRNA-seq and scATAC-seq, provides, to our knowledge, the first comprehensive exploration of photoreceptor developmental differentiation and key regulatory factors in retinoblastoma. Our findings offer potential transcriptional regulatory therapeutic targets and guide the development of novel RB treatment strategies and measures to inhibit RB invasion.

## Methods

### Data extraction

The raw expression matrices of scRNA-seq and scATAC-seq of the retinoblastoma were retrieved from National Center for Biotechnology Information Gene Expression Omnibus under accession number GSE166173 and GSE166174[19].

The raw expression matrices of scRNA-seq and scATAC-seq of the human embryonic developing retina were retrieved from National Center for Biotechnology Information Gene Expression Omnibus under accession number GSE228370[18].

### scRNA-seq preprocessing

We performed alignment to this amended reference using 10x Cellranger (Version 6.0.0),which employs the STAR [55] sequence aligner. The reference genome was the human genome GRCh38. The processed matrices of different samples were merged and the following analyses were finished by Omicverse [56] and Scanpy [57]. We removed cells with less than 200 informative genes expressed, cells with more than 4300 genes expressed and cells with more than 25% of counts corresponding to mitochondrial genes. We removed genes with less than 3 cells expressed. We performed doublet analysis using the ‘Scrublet’ Python package from scanpy [58], and apply doublet filtering to remove doublets biasing. Count data was log-normalized and scaled to 10, 000. We performed high variable gene analysis based parameter setting: ‘min_ mean=0.0125’, ‘max_mean=3’, ‘min_disp=0.5’. Then using PCA embedding to calculate a UMAP layout[59]. Clusters were identified using the Leiden algorithms[60] based on the nearest neighbor graph.

### scATAC-seq preprocessing

Raw sequencing data were converted to fastq format using ‘cellranger-atac mkfastq’ (10x Genomics, v.1.2.0). scATAC-seq reads were aligned to the GRCh38 (hg38) reference genome and quantified using ‘cellranger-atac count’ (10x Genomics, v.1.2.0). Fragment data was further processed using the ‘ChrAccR’ R package (v.dev.0.9.11+). We filtered out cells with less than 1,000 or more than 50,000 sequencing fragments. TSS enrichment was computed as a metric of signal-to-noise ratio using methods described in [61] and we discarded cells with a TSS enrichment less than 4. Fragments on sex chromosomes and mitochondrial DNA were excluded from downstream analysis.

### Data preprocessing for cell paired

To preprocess the scRNA-seq for cell paired, we annotated cell types from scRNA-seq data. To preprocess the scATAC-seq for cell paired, we apply the latent semantic indexing (LSI) for dimension reduction and first encoder transformation. Then we constructed the prior regulatory graph to utilized the multi-omics alignment by GLUE (version 0.3.2)[15].

### Pairing the scRNA-seq and scATAC-seq data by scglue

After constructed the prior regulatory graph from scRNA-seq and scATAC-seq, we specify the probabilistic generative model negative binomial distribution(NB) for the omics-layer to be integrated. Next, we initialize a GLUE model for scRNA-seq and scATAC-seq layers. With the trained GLUE model, eigenvectors for each cell (feature embeddings) were obtained by parameter ‘X_glue’. Subsequently, Pearson coefficients were used to find the two most similar feature vectors in scRNA-seq and scATAC-seq. Duplicate paired cells are discarded in this process.The cell annotations of scRNA-seq were transferred to the corresponding scATAC-seq cells.

### Calculate the ratio of observed to expected cell numbers

The ratio of observed to expected cell numbers (Ro/e) was determined for each cell types cross various samples using the omicverse.utils.roe function with default parameters. This analysis aimed to reveal the preferential distribution of each cell type among the samples. The expected cell numbers of each combination of samples and cell types were obtained from the chi-square test. In brief, we defined that one cell type was identified as being enrichment in a specific sample if Ro/e was greater than 1.5. For visualization, the symbol ‘+++’ denotes instances where the Ro/e exceeds 2, ‘++’ signifies Ro/e greater than 1.5, ‘+’ indicates Ro/e above 1, and ‘+/-’ is utilized to designate Ro/e that is less than or equal to 1.

### Calculation of motif score

We used scbasset (v.0.0.0) [62] to score motifs on a per cell basis using motif injection method. For motif injection, we first generated dinucleotides shuffled background sequences, and inserted motif of interest to the center of those sequences.

### Trajectory inference and pseudotime analysis

Trajectory inference of photoreceptor cells was conducted by using the omicverse.single.pyVIA function in Omicverse. This method is grounded in the VIA Python package, which provides a scalable algorithm for trajectory inference within single-cell RNA sequencing analysis. In addition, trajectory roots were selected based on the RSC score. Furthermore, the pseudotime was depicted as streamlines using the v0.plot_stream function, gene trends were calculated and visualized with the v0.plot_gene_trend function, and the automated detection of terminal states was performed using the v0.plot_trajectory_gams function with default parameters.

### Gene Density Analysis

Gene expression density was computed using the Mellon algorithm[26], which applies a Gaussian kernel to estimate spatial gene distribution based on transcript counts across tissue regions. The algorithm integrates spatial coordinates and normalizes gene expression levels to ensure comparability. Heatmaps were generated to visualize z-score normalized gene densities, highlighting differential expression patterns between tissue regions.

### Differential gene expression analysis

To identify the differentially expressed genes between two groups we specified, we adhered to the ‘Differential Expression Analysis in Single Cell’ tutorial provided by Omicverse on GitHub and employed the omicverse.bulk[56].pyDEG function to evaluate the significance of each gene. The log2 fold change (log2FC) for each gene was determined by calculating the difference between the log2-transformed mean counts of each group. To remove the influence of low expression genes, we selected the top 3000 highly variable genes as the input data. Genes with adjusted P value less than 0.05 (the Wilcoxon rank-sum test) and greater than the threshold of the log2FC were considered as significant differentially expressed genes. Furthermore, the specific threshold for log2FC was clearly indicated in the legend of the respective figures.

### Gene Ontology (GO) and Network-based Analyses of cell subpopulations

We performed gene set enrichment analysis of GO terms using gseapy (v.0.10. 8) [63] with default parameters (adjusted p-values<0.05) and select geneset ‘GO_Biological_Process_2021’ and ‘GO_Molecular_Function_2021’ [64]. We also calculated the fraction of each GO term using ‘Overlap’ based on the results of the enrichment analysis, and logarithmized the adjusted P-value. To distinguish the different Go terms, we transformed the gene of each term into a one-hot code and clustered the terms using ‘clustermap’ of Python package seaborn (v.0.11.2) [65] to find similar term modules. For network-based analyses, protein lists were submitted to the STRIN G-db tool [66] (all active interaction sources, interaction score > 0.8). The network was exported into a simple tabular output, reformatt ed to display desired parameters (e.g., K-means clusters)

### Putative interactions between cell types

To enable a systematic analysis of cell-cell communication, we used CellPhoneDB [67]. CellPhoneDB is a manual curated repository of ligands, receptors and their interactions, integrated with a statistical framework for inferring cell-cell communication networks from single cell transcriptome data. Briefly, in order to identify the most relevant interactions between cell types, we looked for the cell-type specific interactions between ligands and receptors. Only receptors and ligands expressed in more than 10% of the cells in the specific cluster were considered. We performed pairwise comparisons between all cell types. First, we randomly permuted the cluster labels of all cells 1000 times and determined the mean of the average receptor expression level of a cluster and the average ligand expression level of the interacting cluster. For each receptor-ligand pair in each pairwise comparison between two cell types, this generated a null distribution. By calculating the proportion of the means which are as or higher than the actual mean, we obtained a p-value for the likelihood of cell type-specificity of a given receptor-ligand complex. We then prioritized interactions that are highly enriched between cell types based on the number of significant pairs and manually selected biologically relevant ones. For the multi-subunit heteromeric complexes, we required that all subunits of the complex are expressed (using a threshold of 10%), and therefore we used the member of the complex with the minimum average expression to perform the random shuffling.

## Data availability

All data were obtained from publicly available datasets.

## Code availability

All codes used in this manuscript are available in https://github.com/Starlitnightly/Analysis_RB

## Author contribution statement

Y.Z., Z.Z., H.D., and B.Q. designed the experimental workflow. Y.Z., P.L., and L.H. collected the sequencing result data. Y.Z. conducted the main analysis of the manuscript, while P.L. and L.H. jointly analyzed the cellular interactions in Section 4. Y.S. and Y.L. assisted with the analysis of the spatial transcriptomics section in Section 3, and Y.W. assisted with the analysis of transcription factors in Section 3. Y.Z. and P.L. co-wrote the manuscript. Z.Z. validated the results and revised the manuscript.

