## Supplementary material for "Integrated Epigenetic and Transcriptomic Profiling Reveals Dynamic Regulatory Networks Driving Retinoblastoma Pathogenesis"

### Supplement Figure

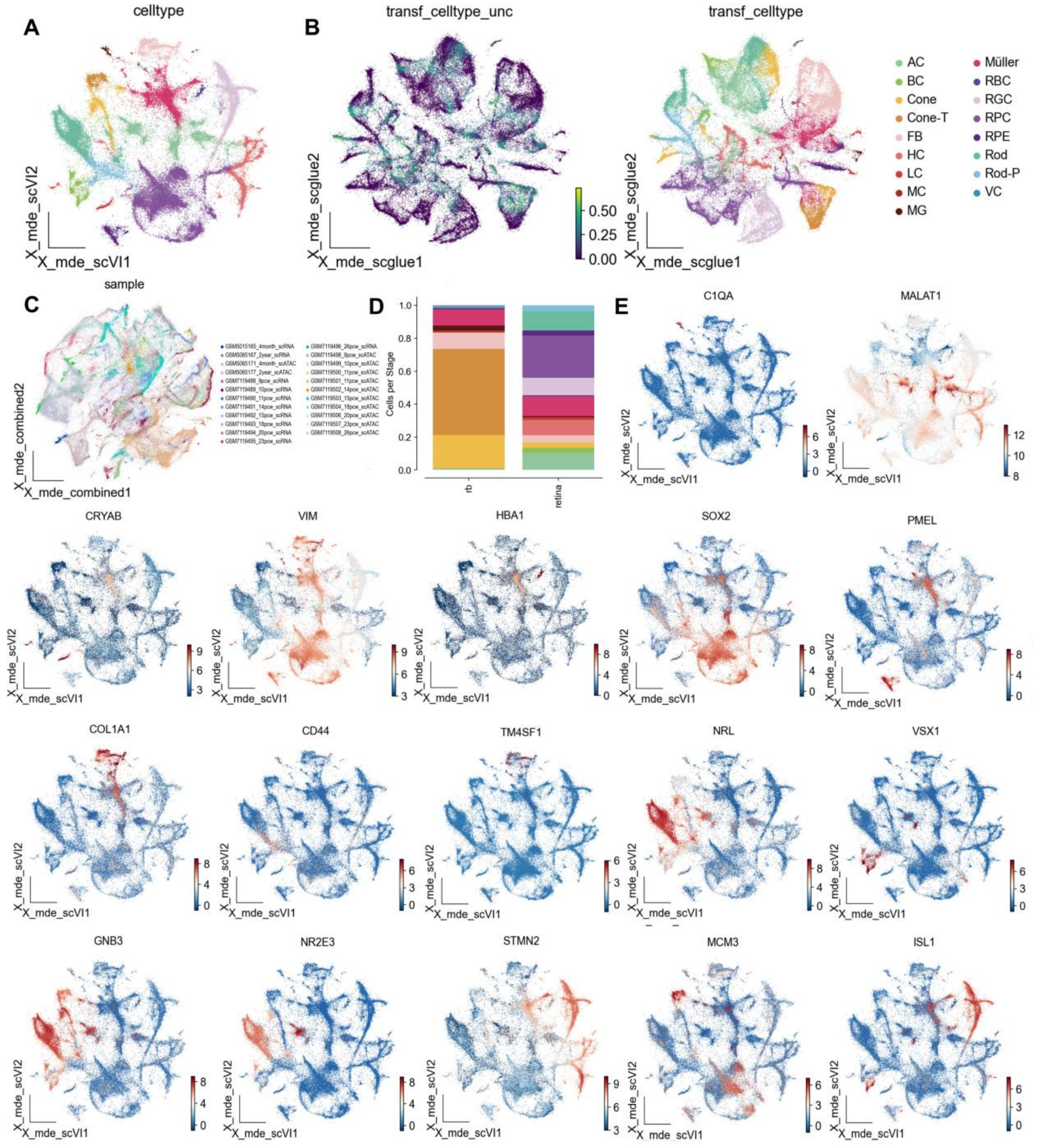


**Supplementary Figure 1. Basic Data Information Atlas of the Retina and Retinoblastoma.**

**A**, UMAP visualization of scRNA-seq data, colored by celltype. **B**, UMAP visualization demonstrates the transfer of scRNA-seq data annotation to scATAC-seq data. **C**, UMAP visualization of integrated multi-omics data, colored by samples. **D**, Bar plots displaying the distribution of cell types across tissues. The shading represents the major cell types. **E**, UMAP visualization illustrating the gene expression patterns of corresponding marker genes, with cells colored according to normalized gene expression levels.

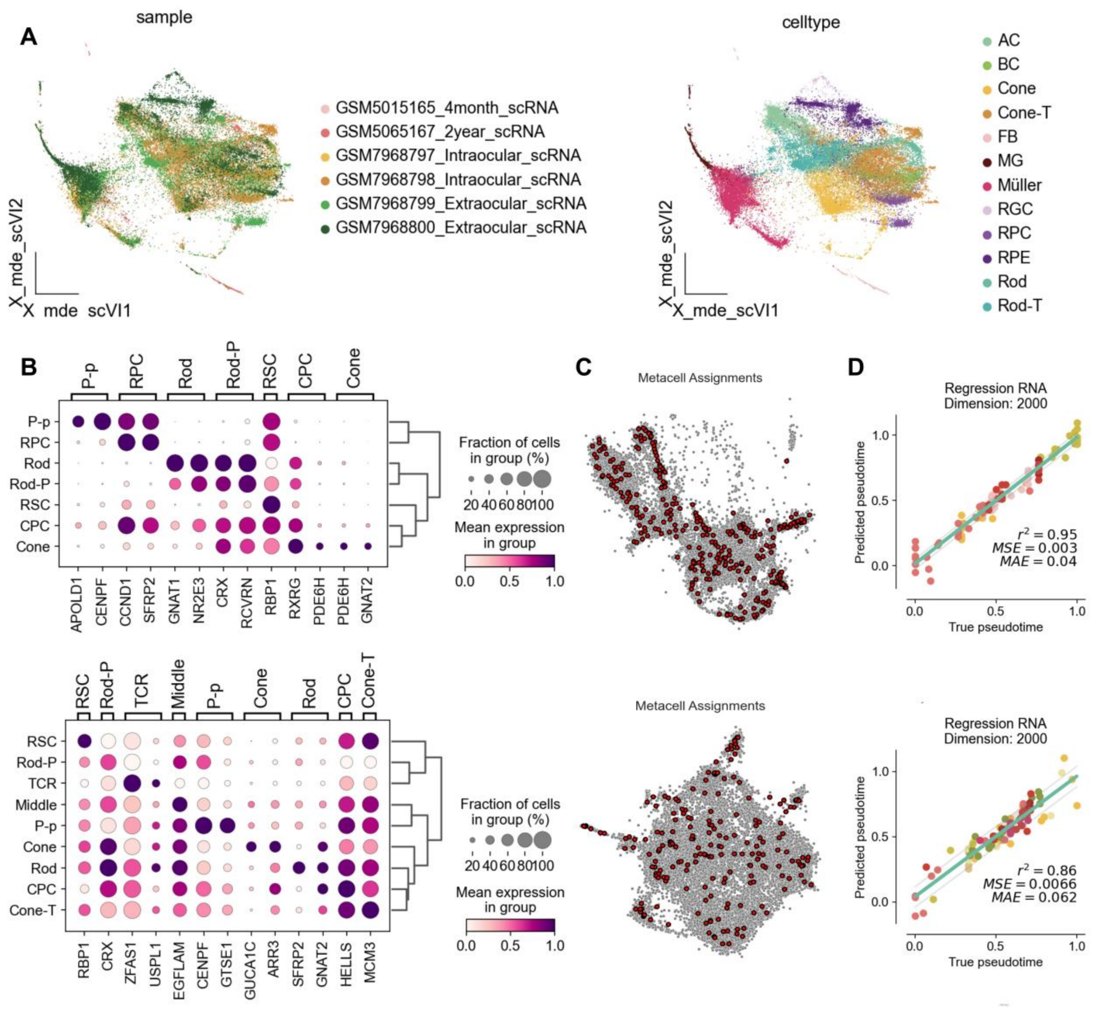


**Supplementary Figure 2. Retinoblastoma overall samples and metacells calculation atlas.**

**A**, UMAP showing the overall sample landscape of retinoblastoma,colored by samples(left) and celltypes(right). **B**, Dot plots displaying the expression patterns of characteristic marker genes in photoreceptor cells. **C**, UMAP showing the calculation and selection of metacells. **D**, A fitted ridge regression model for photoreceptor metacells in the retina (top) and rb (bottom), with the x-axis representing actual pseudotime and the y-axis representing predicted pseudotime.

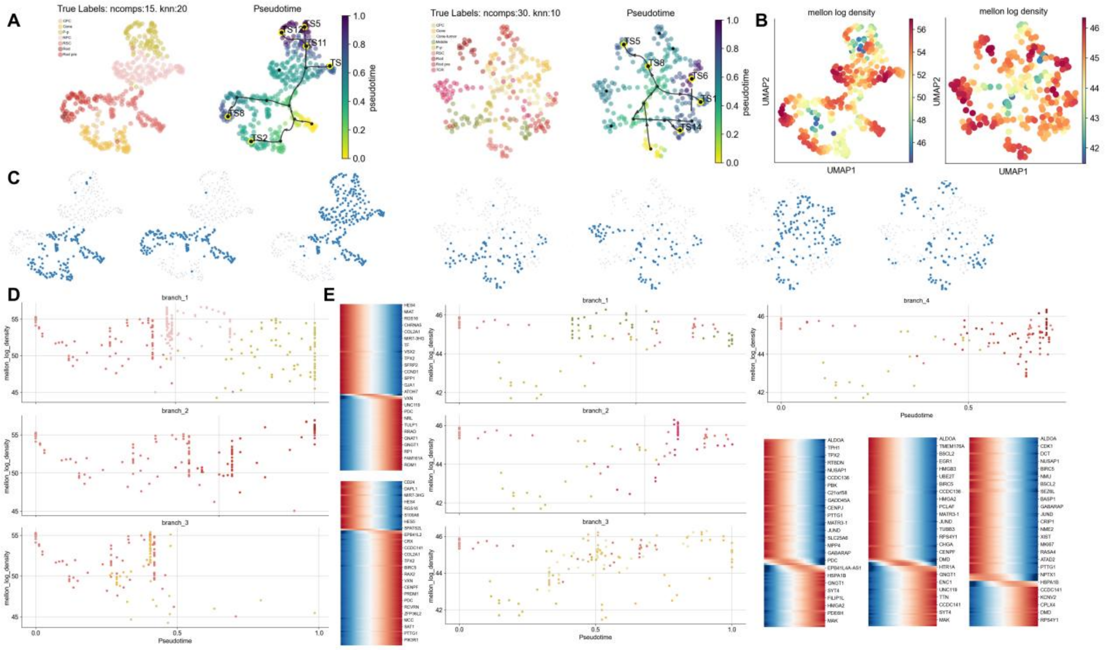


**Supplementary Figure 3. Developmental trajectories and lineages of photoreceptor cells in the retina and retinoblastoma.**

**A**, UMAP showing the pseudotime of photoreceptor metacells.

**B**, Same as A, with UMAPs colored by Mellon log density. Density was computed in high-dimensional cell-state space (diffusion maps).

**C**, UMAPs highlighting the cells spanning RSC to other photoreceptor cells along each lineage.

**D**, Plots comparing Palantir pseudo time to mellon-log-density for different lineages. Top row: RSC→P-p, Middle row: RSC→Rod, Bottom row: RSC→Cone.(left-retina)

Top row: RSC→Middle(left); Rod-T(right), Middle row: RSC→TCR, Bottom row: RSC→Cone-T.(right-rb).

**E**,Heatmaps showing the gene expression dynamics along pseudotime for genes associated with the respective lineage. RSC→Rod (top); Cone (bottom) (left-retina) ; RSC→Middle(left); Rod-T(middle); TCR (right-rb). High and low-density regions were assigned manually by comparing Mellon density with Palantir pseudotime.

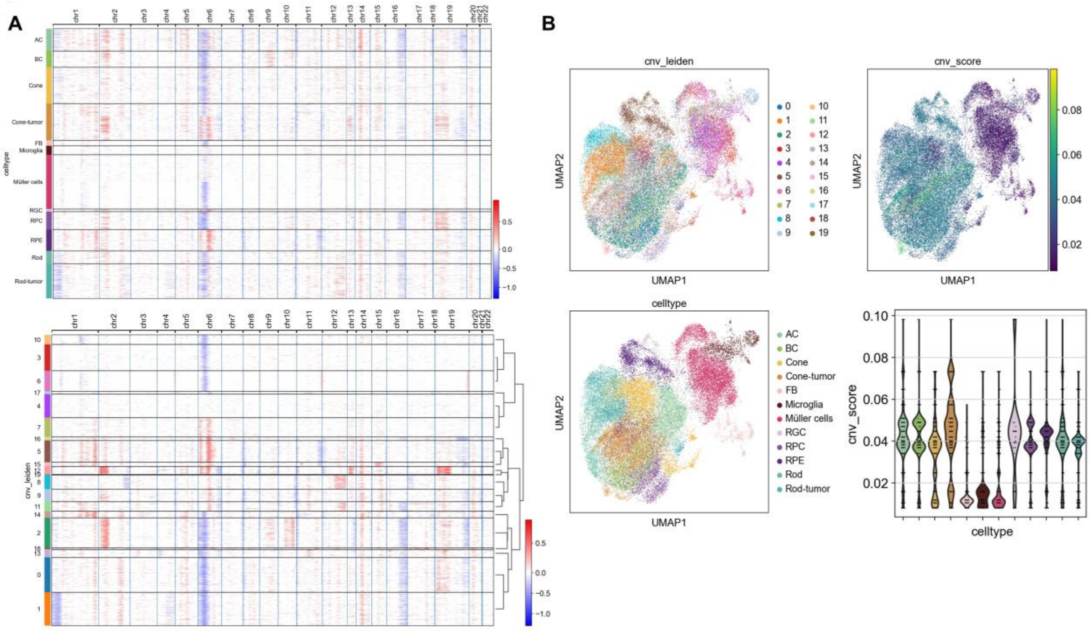


**Supplementary Figure 4. Copy number variation in retinoblastoma data.**

**A**, Copy number variation of distinct cellular subtypes across diverse chromatin domains.

**B**, UMAP and violin plots illustrate copy number variation across different cell subtypes.

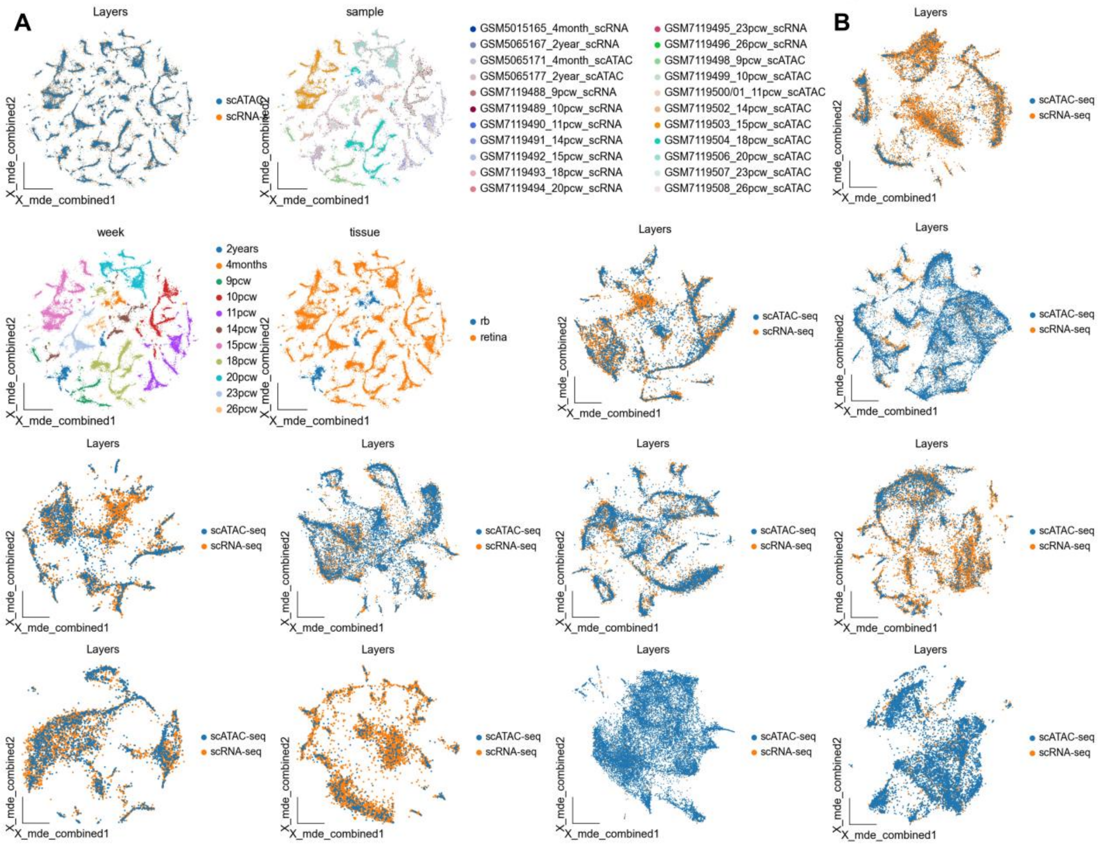

**Supplementary Figure 5. Paired Atlas of scRNA-seq and scATAC-seq Single Samples in Retinoblastoma.**

**A**, UMAP showing the paired scRNA-seq and scATAC-seq single samples in retinoblastoma,colored by layers,samples,week and tissue. **B**, UMAP plot showing the pairwise of single-sample across different weeks (from top to bottom, left to right: 9, 10, 11, 14, 15, 18, 20, 23, 26 pcw, 4 months, 2 years).

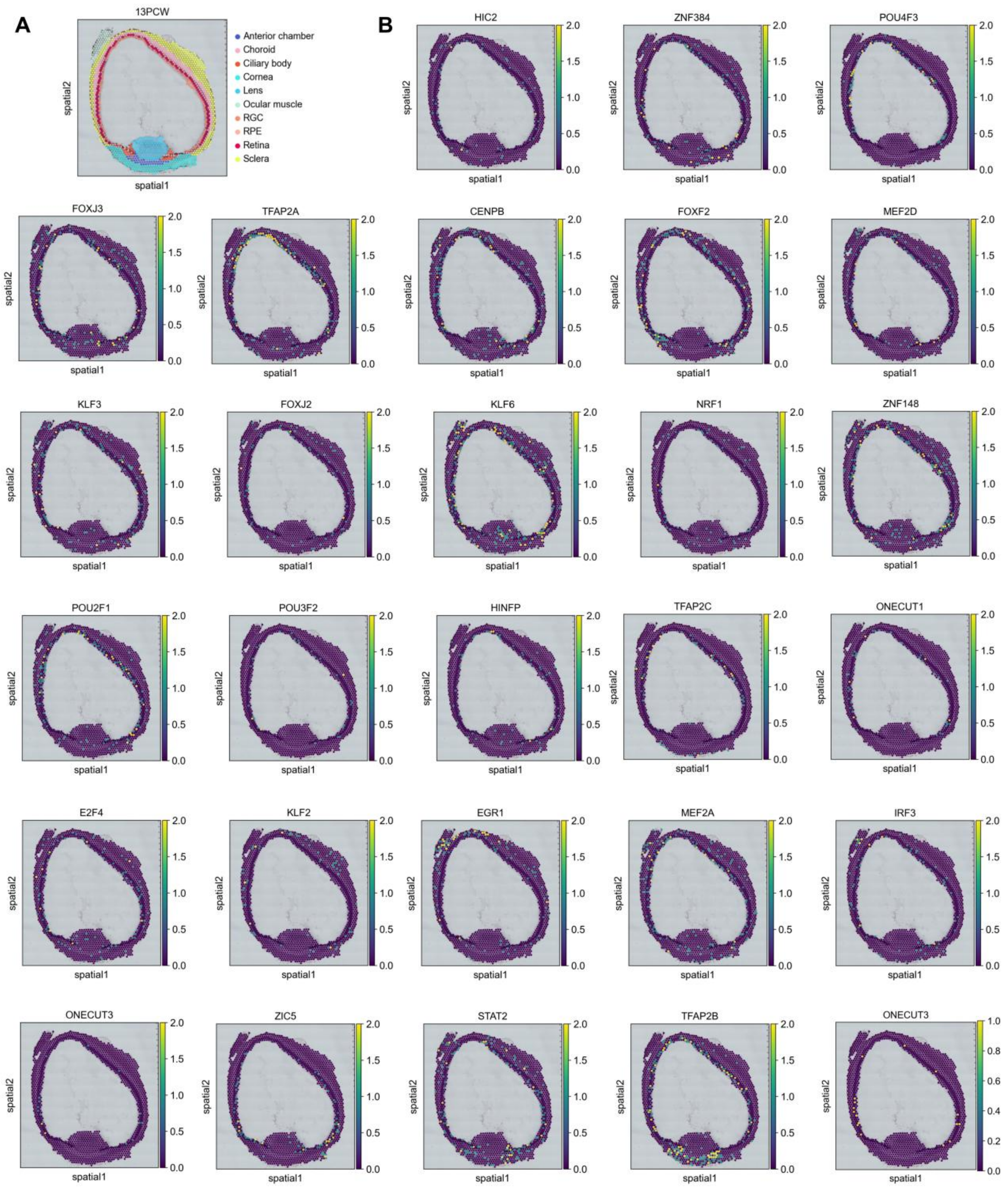


**Supplementary Figure 6. Expression of genes corresponding to highly active TFs in different retinal spatial transcriptomics datasets.** **A**, Annotated schematic of retinal spatial transcriptomics data. **B**, Expression of genes corresponding to some highly active TFs in photoreceptor subtypes within retinal spatial transcriptomics datasets.

### Supplement Table

**Supplementary Table 1.**Top ten most active transcription factors in the P-p, Cone, and Cone-tumor subtypes.

| P-p | Cone | Cone-tumor |
|---|---|---|
| NEUROG2（var.2） | FOXC2 | HINFP |
| NEUROD1 | KLF15 | NRF1 |
| TCF3 | TFAP2B | GRHL1 |
| PTF1A | FOXC1 | ZBTB14 |
| NEUROG2 | FOXJ3 | TFDP1 |
| HAND2 | NRF1 | CTCF |
| TCF21 | BARX2 | KLF15 |
| EBF2 | TFAP2C | E2F6 |
| NEUROD2 | MEF2A | EGR1 |
| ATOH7 | HINFP | TCFL5 |

**Supplementary Table 2.**Regulatory target genes of six specific TFs.

| EBF1 | SOX15 | NFIL3 | ZNF148 | ZBTB14 | YY1 |
|---|---|---|---|---|---|
| PNPLA7 | SDHD | SEC22A | MFAP1 | PHF10 | HNRNPF |
| SLC18A3 | MRPL18 | MRPL55 | SMU1 | HGH1 | COX5B |
| FEM1A | BANP | CHMP1B | KDM5A | WFS1 | CNBP |
| DMPK | HNRNPF | ANXA7 | UTP23 | CAHM | DAD1 |
| OASL | TAF9 | RARA-AS1 | GPATCH8 | XPOT | DNAJC8 |
| FMNL1 | RBMXL1 | LRP5L | DCAF10 | ZNF862 | TPI1 |
| | BCL7C | SRSF8 | YTHDC1 | TBC1D13 | NDUFAB1 |
| | PLEKHH3 | PDSS2 | RBSN | TMEM87B | NDUFC2 |
| | TMEM203 | FAM222A | KMT2B | BMPR1A | CDC37 |
| | MRPL55 | ARRDC3 | AKAP17A | MSANTD2 | TBL1XR1 |
| | LAGE3 | COMT | TBL1XR1 | CTPS2 | OAZ1 |
| | SLC30A4 | LRPAP1 | SETD2 | PIGQ | BNIP3L |
| | PNKD | C14orf119 | FAF2 | NDRG2 | GPX4 |
| | ING2 | RNF13 | DIDO1 | CUX1 | GSTP1 |
| | POLR2C | PPP2R2A | COMMD2 | CERS5 | HNRNPH1 |
| | ACD | AK6 | RAP1A | ZNF646 | AURKAIP1 |
| | C1orf35 | POLR2F | SLK | VPS54 | CTBP2 |
| | TMUB1 | NPTN | UTP6 | | SAFB |
| | AP5S1 | CADM1 | RLF | | EIF4H |
| | DPP7 | ITCH | ELF2 | | STMP1 |
| | GPAA1 | CFL2 | RNF40 | | COX8A |
| | DPH7 | EGLN1 | SMAD4 | | PSMC6 |
| | TCEAL1 | TIMMDC1 | CUL3 | | CHCHD2 |
| | LRPAP1 | CADM4 | NAA15 | | RBM26 |
| | PPAN | BLCAP | RBM6 | | HNRNPA0 |
| | LCMT2 | IL1R1 | DDX42 | | UQCRH |
| | PPP2R2A | ARHGEF1 | PWP1 | | MCRIP1 |

| | | | | | |
|---|---|---|---|---|---|
| | TMEM9 | PTPRH | COPB1 | | RBM25 |
| | THAP7 | SERTAD2 | ZNF236 | | ARL3 |
| | ANKRD39 | OSTM1 | CLK2 | | EDF1 |
| | DBP | RSPRY1 | TJP1 | | SNRNP200 |
| | POLR2F | LINC01679 | PEX1 | | TTC3 |
| | SOX15 | CD3G | TM9SF3 | | COX7A2 |
| | PMF1 | | PRMT2 | | PRDX2 |
| | NDUFV1 | | C15orf40 | | PRPF8 |
| | PMVK | | SETX | | CCT3 |
| | SNHG12 | | HIP1 | | EIF2S2 |
| | AK2 | | ANKRD26 | | NDUFB11 |
| | CADM1 | | NPIPB13 | | THRAP3 |
| | KEAP1 | | | | OASL |
| | SLC39A3 | | | | TMIGD3 |
| | PRPS1 | | | | |
| | SELENOS | | | | |
| | RANGRF | | | | |
| | PLPP1 | | | | |
| | TCEAL3 | | | | |
| | IL1R1 | | | | |
| | OASL | | | | |
| | NPIPB13 | | | | |
| | TMIGD3 | | | | |
